# Sorcin-STAT3-Serpin E1/CCL5 axis can be the trigger of pancreatic cancer-associated new-onset diabetes

**DOI:** 10.1101/2023.07.20.549805

**Authors:** Jiali Gong, Xiawei Li, Zengyu Feng, Jianyao Lou, Kaiyue Pu, Yongji Sun, Sien Hu, Jian Wu, Yulian Wu

## Abstract

A rise in blood glucose is the early warning signs of underlying pancreatic cancer (PC), which could be the externalization of genetic events in PC progression. But there is still a vacancy in the field of mechanism research on pancreatic cancer-associated new-onset diabetes (PCAND). Using siRNA-mediated gene knockdown in vitro, we made MIN6 cells incubated with conditioned media from transfected PC cells, and detected its response. Immunological techniques were used to explore the interaction between sorcin and STAT3. Human cytokine array was performed to explore the inflammatory cytokines downstream of sorcin. In the present study, we have identified a PCAND driver gene SRI. In PC cells, sorcin and STAT3 form a positive feedback loop to enhance the transcription of serpin E1 and CCL5, which can impair nearby islet β-cells, likely by activating the p38 pathway. In 88 biopsies, expression of sorcin was elevated in PC tissues, especially so in PCAND patient samples. Furthermore, clinical-SRI gene combination model can better distinguish PCAND from T2DM, and serpin E1 level is higher in the peripheral blood samples from PCAND than T2DM. Thus, Sorcin could be the key driver in PCAND, and figuring out sorcin-STAT3-serpin E1/CCL5 signaling axis can help us better understand the pathogenesis of PCAND and identify potential biomarkers.

**Statement of significance:** This study mapped out a novel sorcin-STAT3-Serpin E1/CCL5 signaling axis in pancreatic cancer cells, which explains how early pre-symptomatic pancreatic cancer may coincide with new-onset diabetes in some patients.

## INTRODUCTION

Pancreatic cancer (PC) is a highly fatal disease with a 5-year cumulative survival rate of about 10% in the USA [1, 2]. Early diagnosis of PC at a resectable stage opens up more treatment options and substantially improves patient survival[3]. Previous studies of pancreatic tumorigenesis suggested that there is certain timing in the mutation of PC driver genes, activating mutation in *KRAS* are present in low-grade pancreatic intraepithelial neoplasia (PanIN-1) lesions [4] (94.1% mutation rate[5]), and inactivating mutations of *CDKN2A* (17.0%), *TP53* (63.9%) and *SMAD4* (20.8%) were orderly found in the transformation of PanIN-2 and PanIN-3 lesions[6, 7]. However, due to the long time and inadequacy of these gene mutations to PanINs-PDAC evolution[8], related early diagnosis strategies have not achieved significant clinical benefits.

New-onset diabetes, especially in individuals aged over 50 years, has been identified as an early warning sign for underlying PC. A case-control study found that on average patients with pancreatic ductal adenocarcinoma (PDAC) develop hyperglycemia 36 to 30 months before their tumor diagnosis[9], presenting a potential window of opportunity for early detection. Distinguishing new-onset diabetes from the more prevalent type 2 diabetes mellitus (T2DM) is a prerequisite for carrying out targeted screening of this high-risk population. A recent study by Bao *et al.* suggests that pancreatic cancer-associated new-onset diabetes (PCAND) is characterized primarily by reduced insulin secretory capacity resulting from β-cell dysfunctions[10]. Insulin resistance, though also present[11], appears to be less severe than that observed in patients with T2DM[10]. Recent studies have also identified a growing list of biomarkers associated with PCAND[12-23], but the mechanistic link between PC and the pathogenesis of new-onset diabetes remains largely unclear. Actually, the challenge now is how to identify only 1% PCAND from common T2DM in the new-onset diabetes population[24-26].

To identify the key regulator(s) in PCAND pathogenesis, we focused on genetic factors both associated with PC and involved in islet insulin secretion or β-cell dysfunction. *SRI*, which encodes a protein named sorcin (soluble resistance related calcium binding protein), was previously found to act as a β-cell protective factor in T2DM[27]. Interestingly, we found that sorcin was significantly overexpressed in tumor samples from PDAC patients, and especially so in PCAND patient. *In vitro*, we showed that sorcin forms a positive feedback loop with STAT3 and activates the transcription of inflammatory factors, such as CCL5 and serpin E1, to impair nearby pancreatic β-cells. Finally, we preliminarily confirmed the potential of *SRI* and its downstream serpin E1 in distinguishing PCAND from T2DM based on large-scale database and small clinical cohorts.

## MATERIALS AND METHODS

### Cell culture

AsPC-1, PANC-1, CFPAC-1, BxPC-3, Mia Paca-2, HPDE6 and MIN6 cell lines were purchased from American Type Culture Collection (ATCC). PANC-1 and HPDE6 cells were cultured in DMEM (Gibco, USA), and ASPC-1 cells were maintained in RPMI-1640 medium (Gibco, USA). MIN6 cells were cultured in RPMI-1640 medium with 50 μM β-Mercaptoehanol (Cienry, Zhejiang, China). Both DMEM and RPMI-1640 media were supplemented with 10% FBS (YEASEN, Shanghai, China), 100 units/mL of penicillin and streptomycin (Cienry, Zhejiang, China). All cells were cultured in a humidified incubator at 37 °C in a 5% CO_2_ atmosphere. Cells were passaged when 80–90% confluence was reached, and the media were changed every 2 days.

### Plasmid and small interfering RNA (siRNA) transfection

Human pancreatic cancer cells were transfected with plasmids using JetPRIME transfection reagent (Polyplus, Beijing, China). Human pancreatic cancer cells were transfected with siRNA using INTERFERin transfection reagent (Polyplus, Beijing, China) according to the manufacturer’s instructions. The pcDNA-*SRI*-FLAG plasmid was purchased from WZ Biosciences Inc. (Shangdong, China) and pcDNA-*STAT3*-FLAG plasmid was kindly provided by Prof. Hong Zhu (Zhejiang University, Hangzhou, China). The sense strands of the duplex siRNAs (synthesized by GenePharma, Shanghai, China) were: *NC:* 5’-UUCUCCGAACGUGUCACGUTT-3’, *SRI*: 5’-GCCGGCUUAUGGUUUCAAUTT-3’, *STAT3* #1: 5’-GGGACCUGGUGUGAAUUAUTT-3’, *STAT3* #2: 5’-CCCGGAAAUUUAACAUUCUTT-3’, *STAT3* #3: 5’-GGUACAUCAUGGGCUUUAUTT-3’.

### Collection of conditioned media

The culture media of transfected PC cells were changed to RPMI-1640 without FBS at 6 hours after transfection. After 2 days of culturing, the media were collected and centrifuged at 3000 rpm for 15 min at 4°C. The supernatants (conditioned media) were stored at −80 °C and used within one month. In all subsequent experiments, MIN6 cells were incubated with conditioned media supplemented with 5% FBS.

### MTT assay for cell viability

MIN6 cells were seeded in 96-well plates (5000 cells/well) and incubated in different conditioned media for 72 hours, with medium change every 24 hours. Then 20 μL of MTT stock solution was added to the culture medium. After incubating at 37 °C for 4 hours, the medium was completely removed, and 200 μL of DMSO were added to the cells to dissolve the violet crystals. Optical density (OD) values were measured using a microplate reader.

### Apoptosis Detection

MIN6 cells were seeded in 12-well plates (1×10^5^ cells/well) and incubated in different conditioned media for 72 hours, with medium change every 24 hours. The cells were then harvested by trypsin digestion. Apoptotic cells were stained using the Annexin V-PI apoptosis detection kit (Yeasen, Shanghai, China) and immediately analyzed by flow cytometry.

### Western blot analysis

Cells were lysed with RIPA lysis buffer (Sigma-Aldrich, USA) supplemented with a protease inhibitor mixture (Thermo, USA), and proteins were quantified using the BCA assay. The experimental procedures were performed in accordance with standard protocols. Briefly, proteins were separated on 15% SDS-polyacrylamide gels and blotted onto polyvinylidene difluoride (PVDF) membranes. Membranes were blocked with 5% milk, incubated with specific primary antibodies, horseradish peroxidase (HRP)-conjugated secondary antibodies, and subsequently subjected to chemiluminescence detection. The antibodies used and the corresponding dilutions are listed in Supplementary Table 1.

### Quantitative real-time PCR (qRT-PCR)

Gene expression levels were assessed using qRT-PCR. Total RNA was extracted from PC cells and MIN6 cells with TRIzol reagent (Invitrogen, USA). The cDNA templates were synthesized with PrimeScript RT Reagent Kit (TaKaRa, China), and qRT-PCR was performed with a 7500 Fast™ System (Applied Biosystems, USA). *GAPDH* was chosen as an endogenous control. Data were analyzed using the 2^−ΔΔCt^ method. Specific miRNA primers are listed in Supplementary Table 2.

### Immunofluorescence and confocal microscopy

Immunofluorescence analysis was performed to measure insulin levels in MIN6 cells. After 72 h of culturing with different conditioned medium, the cells were washed and fixed in prechilled methanol for 10 min and then permeabilized with 0.1% Triton X-100 for another 10 min. After blocking with 5% BSA, the cells were incubated overnight at 4 ℃ with an anti-insulin Ab. The primary Ab was detected by cy3-conjugated goat anti-rabbit IgG goat polyclonal antibody for 1 h (HUABIO, Hangzhou). Cell nuclei were stained with DAPI. Observation and image acquisition were performed using a confocal microscope (Zeiss, Germany). The antibodies used and the corresponding dilutions are listed in Supplementary Table 1.

### Immunohistochemistry and scoring standards

Immunohistochemical analysis was performed to assess sorcin, p-STAT3 and insulin expression levels in these samples. Paraffin-embedded tissues were cut into 4 μm sections and deparaffinized. The sections were incubated with anti-sorcin, anti-p-STAT3 and anti-insulin antibodies respectively overnight at 4 ℃. The antibodies used and the corresponding dilutions are listed in Supplementary Table 1. After incubation with biotinylated goat anti-mouse IgG for 30 min at 37 ℃, each slide was rinsed in phosphate-buffered saline and incubated in the streptavidin-biotinylated horseradish peroxidase complex for 30 min at 37 ℃. The slides were developed using diaminobenzidine substrate solution to visualize peroxidase signals and then counterstained with hematoxylin. These immunohistochemical stains were evaluated by three experienced pathologists independently in a blinded manner, and graded on a semiquantitative scale according to the percentage (P) of stained area (0% area stained = 0, 1–25% area stained = 1, 26–50% area stained = 2, and 51–100% area stained = 3) and the intensity (I) of staining (no staining = 0, weak staining = 1, moderate staining = 2, and strong staining = 3). Results were scored by multiplying the percentage of stained area by the intensity (P × I; maximum = 12). All cases were divided into two groups: a high expression group (with a score ranging from 7 to 12) and a low expression group (with a score ranging from 0 to 6).

### Candidate gene screening process

A total of 213 human genes associated with the Gene Ontology term “insulin secretion” (GO:0030073) were retrieved from the MSigDB database (time of download: 2021/3). The screening process was performed using the Gene Expression Profiling Interactive Analysis (GEPIA) website (http://gepia.cancer-pku.cn/). We first compared the expression levels of each gene in PC tissues vs. adjacent normal tissues. Those genes showing significantly higher expression in PC tissues were further subject to survival analysis, where the association between gene expression and prognosis were tested by comparing the overall survival and disease-free survival for patients in the high expression group (gene expression level ≥ population median) vs. those in the low expression group (gene expression level < population median).

### Glucose-stimulated insulin release assay (GSIS)

To set up the GSIS assay, MIN6 cells were seeded in 12-well plates (1×10^5^ cells/well) and incubated in different conditioned media for 72 hours, with media change every 24 hours. For the GSIS assay, the cells were pre-incubated in Krebs-Ringer bicarbonate buffer (KRBB: 119 mM NaCl, 2.5 mM CaCl_2_, 1.19 mM KH_2_PO_4_, 1.19 mM Mg_2_SO_4_, 10 mM HEPES [pH 7.4], and 1% bovine serum albumin) for 1 hour at 37 ℃, then switched to KRBB containing 5.6 mM of glucose (“low-Glu”) for 1 hour and then 16.7 mM of glucose (“high-Glu”) for another hour. The buffers were collected separately, and centrifuged at 1500 rpm for 15 min. All supernatants were stored at −80 °C and used within 2 weeks. The amount of insulin released into the KRBB buffer were determined using an ELISA kit (CUSABIO, Wuhan).

### Human cytokine array

A membrane-based antibody array (Proteome Profiler Human Cytokine Array Kit, R&D Systems, ARY005B) was used to profile 36 soluble proteins, mostly cytokines and chemokines, in the conditioned medium from PANC-1 cells transfected with either *SRI*-siRNA or NC-siRNA. The complete list of proteins represented in this antibody array can be found at the manufacturer’s website (https://www.rndsystems.com/products/proteome-profiler-human-cytokine-array-kit_ary005b).

### Clinical study design and population

88 pancreatic ductal adenocarcinoma (PDAC) biopsies, consisting of 32 patients without diabetes (pure PC), 28 patients with new-onset diabetes (PCAND, with diabetes diagnosed 24 months before diagnosis of PDAC[28]), and 28 patients with long-standing T2DM (PC+T2DM, with diabetes diagnosed > 24 months before the diagnosis of PDAC) who were enrolled at the Second Affiliated Hospital of Zhejiang University between 2013 January and 2017 December. The diagnostic criteria for T2DM was in accordance with the American Diabetes Association[29]. All pancreatic biopsies were classified according to the American Joint Committee on Cancer (AJCC) Staging Manual, 6th Edition. 21 peripheral blood samples, consisting of 8 PCAND cases and 13 T2DM cases, were collected between 2018 January and 2021 January. This study was approved by the ethics committees of Zhejiang University.

### Measurement of cytokine levels

Peripheral blood samples from pancreatic cancer with new onset diabetes (n = 8) and type 2 diabetes patients (n = 13) were collected between January 2018 and January 2021 at the Second Affiliated Hospital of Zhejiang University. After anticoagulant treatment and centrifugation at 3000 rpm for 10 min, plasma concentrations of the cytokines CCL5 and serpin E1 were measured with an ELISA kit (CUSABIO, Wuhan, China) according to the manufacturer’s instructions.

### UK Biobank Database Study Design and population

The UK Biobank is an ongoing project to demonstrate the successful collection and sharing of linked genetic, physical and clinical information on a population scale. Extensive genetic and clinical data have been collected for approximately 500,000 volunteers across the United Kingdom[30]. We identified patients with cancer using the International Classification of Diseases codes (versions 10) that were recorded by the national cancer registry based on hospital admissions and cause of death. T2DM cases were defined as having an ICD-10 code of E11.X. Only cases in which the individuals did not have T2DM and cancer at the date of attending assessment center were included in this research and subsequently followed up for incident T2DM and PDAC events. Then, participants were split into groups of PCAND or T2DM according to the following criteria: PCAND if diagnosed with PDAC within 24 months after diagnosis of T2DM; T2DM if no cancer occurred during the follow-up which was longer than 36 months after diagnosis of T2DM. The current research was conducted using the UK Biobank Resource under Application 91799.

### Model construction

The information of single nucleotide polymorphisms (SNPs) located in *SRI* and other oncogenes are from National Library of Medicine (https://www.ncbi.nlm.nih.gov/snp). Univariate logistic regression was used to select the significant SNPs that could be used to distinguish PCAND from T2DM. Significant SNPs were identified to be integrated into the combined models. 76 unique clinical variables recorded in the UK Biobank were used to build clinical risk model. The performance of the final logistic regression-based models was assessed in the validation set.

### Model construction

In order to build models to differentiate PCAND from T2DM, the cohort from the UK Biobank database was randomly split (8:2) into the training and testing sets, using the Python (version 3.9.0) package scikit-learn.

76 unique clinical variables in total were collected including 10 physical measures indicators (pulse rate, diastolic blood pressure, systolic blood pressure, waist circumference, weight, body mass index (BMI), hip circumference, standing height, seated height, sitting height) and all variables in the categories of blood count and blood biochemistry during the first assessment visit period (2006–2010), each of which was assessed by logistic regression in the training set while adjusting for age and sex. Indicators with missing data in more than 10% of the population or with no significance (*P* ≥ 0.05) were removed and finally only 9 of them (Standing height; Sitting height; Alkaline phosphatase; Glycated haemoglobin (HbA1c); Total bilirubin; Haemoglobin concentration; Mean corpuscular haemoglobin; Mean corpuscular haemoglobin concentration; Reticulocyte count) were used to build clinical risk model.

Then, we searched the keyword “SRI” in National Library of Medicine (https://www.ncbi.nlm.nih.gov/snp) and got 8462 single nucleotide polymorphisms (SNPs) located in *SRI*, 5 out of which were available in the UKB genotyping data. Univariate logistic regression was used to select the significant variants that could be used to distinguish PCAND from T2DM. Likewise, we analyzed *STAT3*, *KRAS*, *CDKN2A*, *SMAD4* and *TP53* accordingly. Finally, 3 significant variants (rs6465133 (*SRI*) (*P* = 0.001), rs61759623 (*KRAS*) (P = 0.005) and rs104894097 (*CDKN2A*) (P = 0.001)) were identified to be integrated into the combined models, including “clinical+*SRI*”, “clinical+*KRAS*” and “clinical+*CDKN2A*”.

The performance of the final logistic regression-based models was assessed in the validation set. The evaluation indicators used to compare the performance of models were area under the receiver operating characteristic curve (AUROC), sensitivity, specificity and accuracy.

### Statistical analysis

Data were acquired from at least three independent experiments and presented as mean ± SD. All statistical analyses were performed in GraphPad Prism version 8.0.2. Unpaired Student’s t-test was used for comparison between two groups and one-way ANOVA for multiple groups. Kaplan-Meier curves of overall survival were compared using the log-rank test. Correlation coefficients were calculated using the Pearson method. *P* < 0.05 was considered statistically significant.

## RESULTS

### *SRI* is highly expressed in pancreatic cancer and associated with malignant biological behavior

Mounting evidence suggests that PCAND is a paraneoplastic phenomenon caused by paracrine factors secreted by cancer or stroma cells[31, 32], some of which have been shown to impinge on β-cells and inhibit insulin secretion [13, 20, 33]. Finding the common denominator in PC and insulin secretion could help unlock the mechanism of PCAND. Following a screening scheme aimed at finding PC-associated genetic factors involved in insulin secretion (Fig. 1A), we identified *SRI* as the only candidate gene. TCGA and GTEx data showed that sorcin expression was up-regulated in pancreatic adenocarcinoma (PAAD) samples compared with normal pancreatic tissues (Fig. 1B). There was also a positive correlation between high sorcin expression and advanced TNM stages of pancreatic cancer (Fig. 1C). Moreover, elevated expression of sorcin in PAAD was associated with poor overall survival and disease-free survival (Fig. 1D-E).

**Figure 1.**
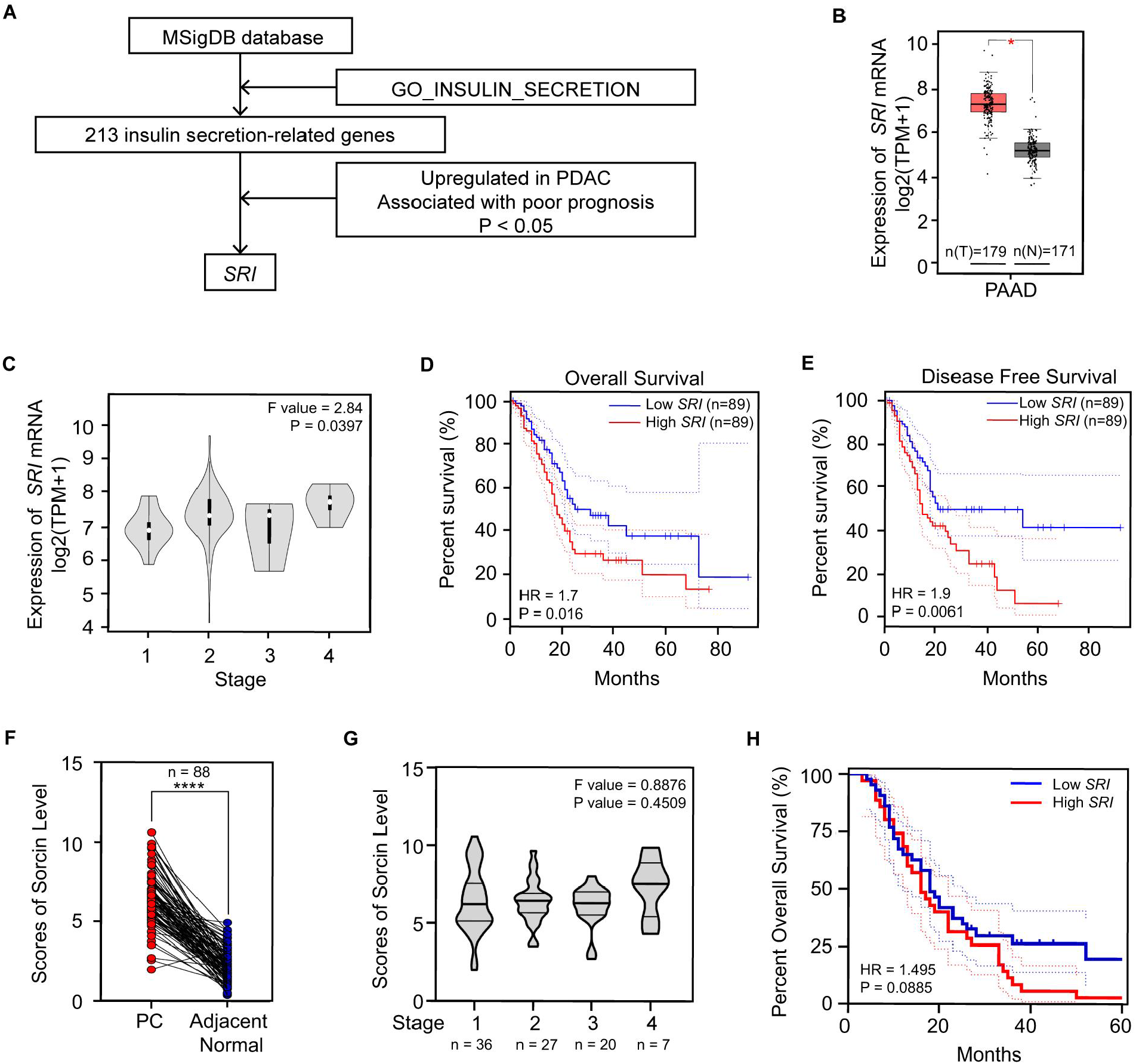
*SRI* is highly expressed in pancreatic cancer and associated with malignant biological behavior. **(A)** The screening process for candidate gene. **(B)** The expression pattern of *SRI* in PC tumor (T) compare with normal pancreas tissue (N). **(C)** The expression pattern of *SRI* in PC patients with different pathological TNM stages. Comparison of **(D)** overall survival rates and **(E)** disease free survival rates between high *SRI* group and low *SRI* group, taking the median of *SRI* expression as the cut-off. **B-E** are based on The Cancer Genome Atlas (TCGA) database and Genotype-Tissue Expression (GTEx). **(F)** Statistical analysis of sorcin level in pancreatic cancer tissues and adjacent normal tissues from 88 patients diagnosed with PDAC. **(G)** Statistical analysis of sorcin level in pancreatic cancer tissues from 88 PDAC patients diagnosed with different pathological TNM stages. **(H)** Comparison of overall survival rates between high *SRI* group (with a score ranging from 7 to 12) and low *SRI* group (with a score ranging from 0 to 6). **F-H** are based on PDAC patients enrolled at the Second Affiliated Hospital of Zhejiang University between 2013 January and 2017 December. *P<0.05; ****P<0.0001, means ± SD was shown. Statistical analysis was performed by Student’s t-test analysis for two groups and one-way ANOVA for multiple groups. And Kaplan-Meier curves of survival were compared using the log-rank test.

To validate these findings in an independent cohort, we examined biopsies obtained from 88 patients diagnosed with pancreatic ductal adenocarcinoma (PDAC). Sorcin expression was significantly up-regulated in tumor tissues compared with adjacent normal tissues (scores of sorcin levels in IHC, PC tissues vs. paired adjacent normal tissues: 6.49 ± 1.68 vs. 2.18 ± 1.02 (n = 88), *P* < 0.0001) (Fig. 1F). However, the expression levels of sorcin were similar in tumor samples of different TNM stages in our cohort (Fig. 1G), probably because TNM stage is a macroscopic and anatomy-dependent system which may not reflect the cancerous behavior of pancreatic cancer. Otherwise, when the patients were categorized based on the amount of sorcin expression detected in their tumor samples, the high *SRI* group and the low *SRI* group had similar median survival times (16 months in the high *SRI* group vs. 18 months in the low *SRI* group), although the rate of death did appear to slow down for patients in the low *SRI* group once they reached 30 months after surgery (Fig. 1H).

### *SRI* is highly expressed in pancreatic cancer-associated new-onset diabetes

Patients in the PDAC cohort can be further classified into three groups based on their diabetes status: those with no diabetes (pure PC), those with new-onset diabetes (PCAND), and those with long-term diabetes (PC+T2DM). Notably, a higher level of sorcin expression was detected in PCAND tumor tissues than in PC+T2DM tumor tissues (scores of sorcin levels in IHC, PCAND vs. PC+T2DM: 7.10 ± 1.71 (n = 28) vs. 5.85 ± 1.67 (n = 28), *P* = 0.008; pure PC vs. PCAND: 6.51 ± 1.51 (n = 32) vs. 7.10 ± 1.71 (n = 28), *P* = 0.136) (Fig. 2A-B). The area under the curve (AUC) for sorcin in differentiating between PCAND patients and PC+T2DM patients was 0.675 (*P* = 0.02423, 95% CI 0.5358-0.8150) (Fig. 2C). Furthermore, fasting blood glucose level in patients with pure PC and PCAND before pancreatectomy was positively correlated with sorcin expression level (Pearson correlation coefficient between scores of sorcin levels in IHC and fasting blood glucose level *r* = 0.281, *P* = 0.0326) (Fig. 2D), which was not observed in PC+T2DM patients (Pearson correlation coefficient between scores of sorcin levels in IHC and fasting blood glucose level, *r* = -0.0572, *P* = 0.7722) (Fig. 2E), suggesting a potential link between the up-regulation of sorcin and islet dysfunction specific to PCAND. Interestingly, in rare instances where the pancreatic tissue section captured both the PDAC tumor and the adjacent islets, high sorcin level in PDAC tumors coincided with low insulin levels in tumor-adjacent islets (Fig. S1). However, since this phenomenon was observed in rare two sections, the conclusion may not be solid and only partially suggested that decreased insulin secretion was likely responsible for causing the higher fasting blood glucose level in patients with high sorcin expression.

**Figure 2.**
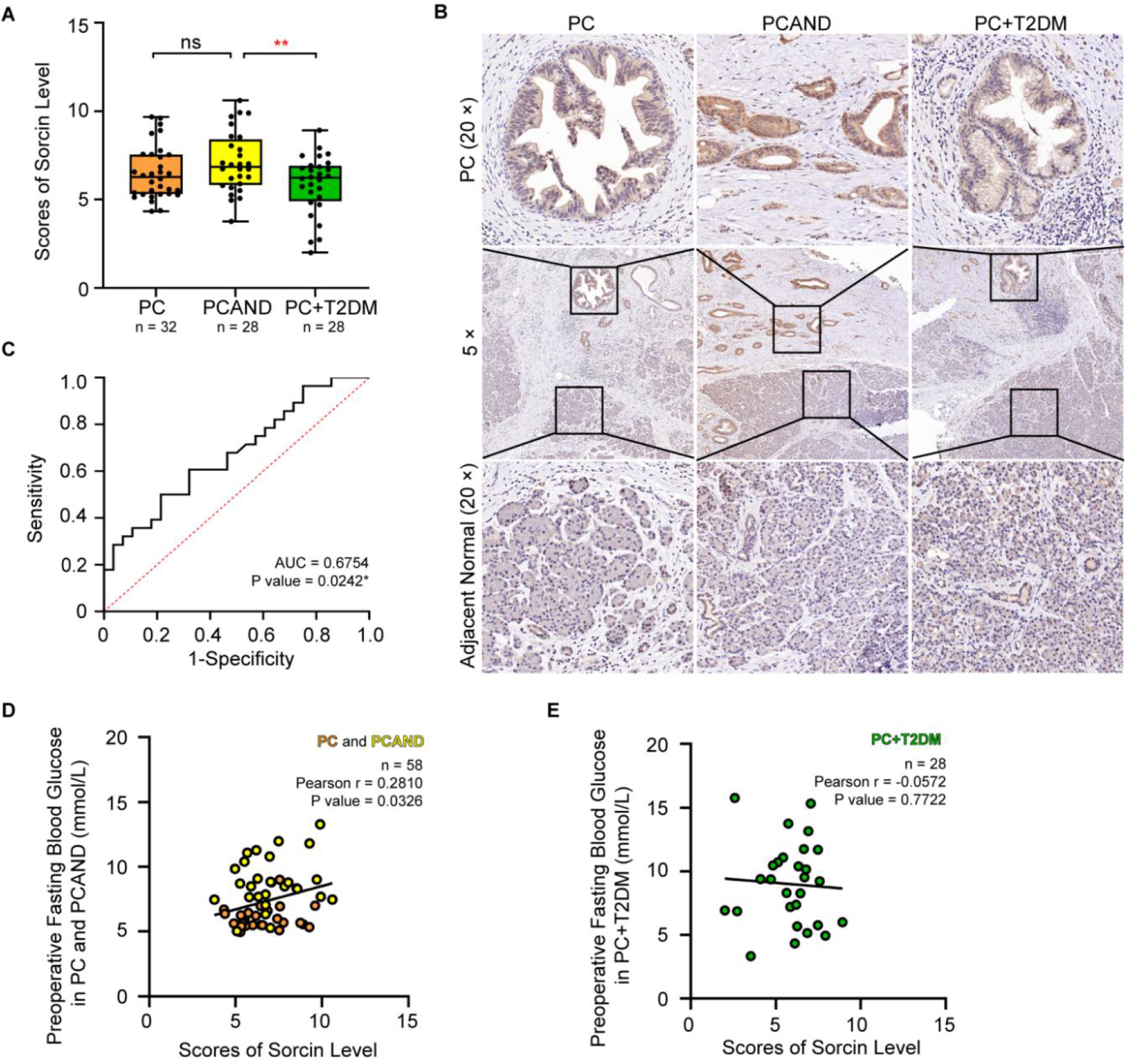
SRI is highly expressed in pancreatic cancer-associated new-onset diabetes. **(A)** Statistical analysis of sorcin level in pancreatic cancer tissues from patients with pure pancreatic cancer (PC), pancreatic cancer with new onset diabetes (PCAND) and pancreatic cancer with long-term diabetes (PC+T2DM). **(B)** Representative images of sorcin staining in tumor tissues and adjacent normal tissues form patients with PC, PCAND and PC+T2DM. **(C)** The sensitivity and specificity in discriminating PCAND patients from PC+T2DM population for sorcin are shown. The ROC curves and AUC were calculated in 28 PCADN patients and 28 PC+T2DM patients. Correlation analysis of fasting glucose level before pancreatic tumor resection and sorcin immunohistochemical score in **(D)** PDAC patients without T2DM (pure PC and PCAND, green point) and **(E)** with T2DM (PC+T2DM, orange point). Ns, no significance; **P<0.01, means ± SD was shown. Statistical analysis was performed by Student’s t-test analysis for two groups. Correlation analysis was performed using the Pearson method on linear regression analysis.

### PC cells inhibit insulin secretion in MIN6 cells in a sorcin-dependent manner

To investigate how sorcin up-regulation may lead to islet dysfunction in PCAND, we utilized *in vitro* cell cultures to mimic the interactions between pancreatic cancer and islet tissue (Fig. 3A). All five PC cell lines we tested (PANC-1, CFPAC-1, BxPC-3, Mia Paca-2, and AsPC-1) recapitulated the elevated expression of sorcin found in patient tumor samples, when compared with a normal pancreatic duct cell line HPDE6 (Fig. 3B). The same expression patterns were observed in two published external datasets, GSE138437 and GSE166165 (Fig. S2A-B). Previous studies have established that insulin-secreting cell lines, such as MIN6 and INS-1, exhibit impaired glucose-stimulated insulin secretion (GSIS) when co-cultured with PC cells or treated with conditioned media from PC cells[13, 20, 33]. To test whether high expression of sorcin is required for this process, we performed a knockdown experiment using small-interfering RNAs (siRNAs). Three PC cell lines with particularly high sorcin expression, PANC-1, AsPC-1 and CFPAC-1, were transfected with either siRNA against *SRI* (*SRI*-siRNA) or negative-control siRNA (NC-siRNA). The knockdown efficiency of *SRI*-siRNA was estimated to be about 15%-40% by Western blot (Fig. 3C and S2C). MIN6 cells exposed to conditioned media collected from NC-siRNA-transfected PC cells (CM-NC-siRNA) showed suppressed GSIS response (Fig. 3D and S2D-E), decreased insulin content (Fig. 3E-G and S2F-H) and decreased expression of transcripts genes related to insulin synthesis and secretion [34-38] (Fig. 3H-K and S2I-L), as quantified by qRT-PCR. This phenomenon was accompanied by lower viability (Figs. 3L-M and S2M-N) and enhanced apoptosis in MIN6 cells (Fig. 3N-O and S2O-P). The deleterious effects of conditioned media on MIN6 cells were partially rescued when the expression of sorcin in the PC cells was knocked-down by *SRI*-siRNA (CM-*SRI*-siRNA in Fig. 3C-O and S2C-P). Together, these results suggest that sorcin is involved in the production of paracrine factors by PC cells, which can negatively impact MIN6 cells’ viability and ability to synthesize and release insulin.

**Figure 3.**
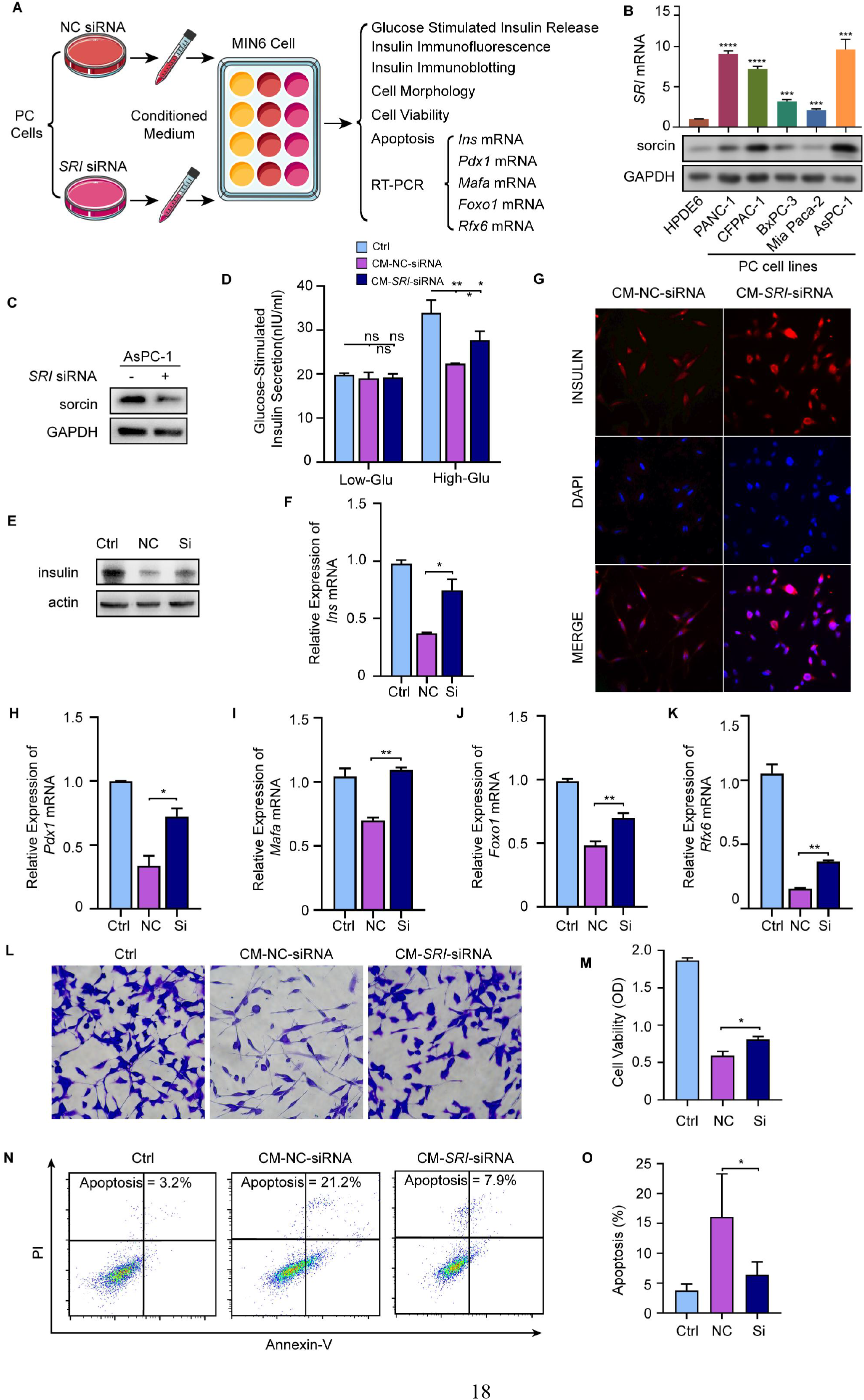
PC cells inhibit insulin secretion in MIN6 cells in a sorcin-dependent manne. **(A)** Schematic diagram of experimental design. **(B)** The mRNA (upper) and protein (lower) levels of *SRI* in PC cell lines and normal pancreatic duct cell line (HPDE6). **(C)** Detection of knockdown effect in ASPC-1 cells by immunoblotting. **(D)** Detection of insulin content in supernatant after GSIS in MIN6 cells incubated with conditioned medium form AsPC-1 cells pretreated with NC siRNA (CM-NC-siRNA) and *SRI* siRNA (CM-*SRI*-siRNA). **(E)** Detection of insulin content in MIN6 cells incubated with CM-NC-siRNA and CM-*SRI*-siRNA from AsPC-1 cells by immunoblotting. **(F)** The expression of *Ins* mRNA in MIN6 cells incubated with different conditioned medium from AsPC-1. **(G)** Immunofluorescence shows the content of insulin in MIN6 cells treated by different conditioned medium from AsPC-1. The expression of **(H)** *Pdx1*, **(I)** *Mafa*, **(J)** *Foxo1* and **(K)** *Rfx6* mRNA in MIN6 cells incubated with different conditioned medium from AsPC-1. **(L)** Morphology of MIN6 cells treated with different conditioned medium from AsPC-1. **(M)** Detection of cell viability of MIN6 cells treated with different conditioned medium from AsPC-1 by MTT assays. **(N)** Detection of apoptosis by flow cytometry in MIN6 treated with different conditioned medium from AsPC-1. **(O)** Quantification results of apoptosis rate. Ns, no significance; *P<0.05; **P<0.01; ***P<0.001; ****P<0.0001, means ± SD was shown. Statistical analysis was performed by Student’s t-test analysis for two groups.

### Sorcin-overexpressing PC cells release CCL5 and serpin E1 to inhibit insulin secretion in MIN6 cells

To further elucidate the mechanism by which conditioned media from PC cells impact MIN6 cells, we set out to identify the paracrine factors released by PC cells under the regulation of sorcin. The pancreatic tumor microenvironment is known to be rich in inflammatory cytokines that support tumor growth[39, 40] and contribute to β-cell dysfunction and apoptosis[41]. To test the possibility that sorcin-overexpressing PC cells release inflammatory cytokines, we used a human cytokine array to analyze the cytokine profile in the supernatants (conditioned media) collected from PANC-1 cells with and without sorcin knockdown. Five cytokines were found to be significantly down-regulated in the *SRI*-siRNA group (Fig. 4A-B). Among them, only CCL5 and serpin E1 were consistently down-regulated in all five PC cell lines following sorcin knockdown (Fig. 4C and S3A-D). Recombinant CCL5 and serpin E1 proteins each inhibited the GSIS response in MIN6 cells in a dose-dependent manner (Figs. 4D-E), confirming the role of these inflammatory cytokines in disrupting β-cell insulin secretion. Moreover, treatment with CCL5 and serpin E1 for prolonged durations (> 48h for CCL5 and > 12h for serpin E1) led to an increase in p38 mitogen-activated protein kinase (MAPK) activation in MIN6 cells (Fig. 4F-G), which has been associated with β-cell apoptosis[42] and may also underlie the apoptotic phenotype induced by conditioned media from sorcin-overexpressing PC cells (CM-NC-siRNA in Figs. 3L-O and 4H). Remarkably, the PC-induced p38 activation in MIN6 cells was attenuated when sorcin expression was knocked down (CM-*SRI*-siRNA in Fig. 4H). Thus, we have identified CCL5 and serpin E1 as key components of PC cell secretions that disrupt β-cell functions, and p38 as a potential downstream target of these inflammatory cytokines in β-cells.

**Figure 4.**
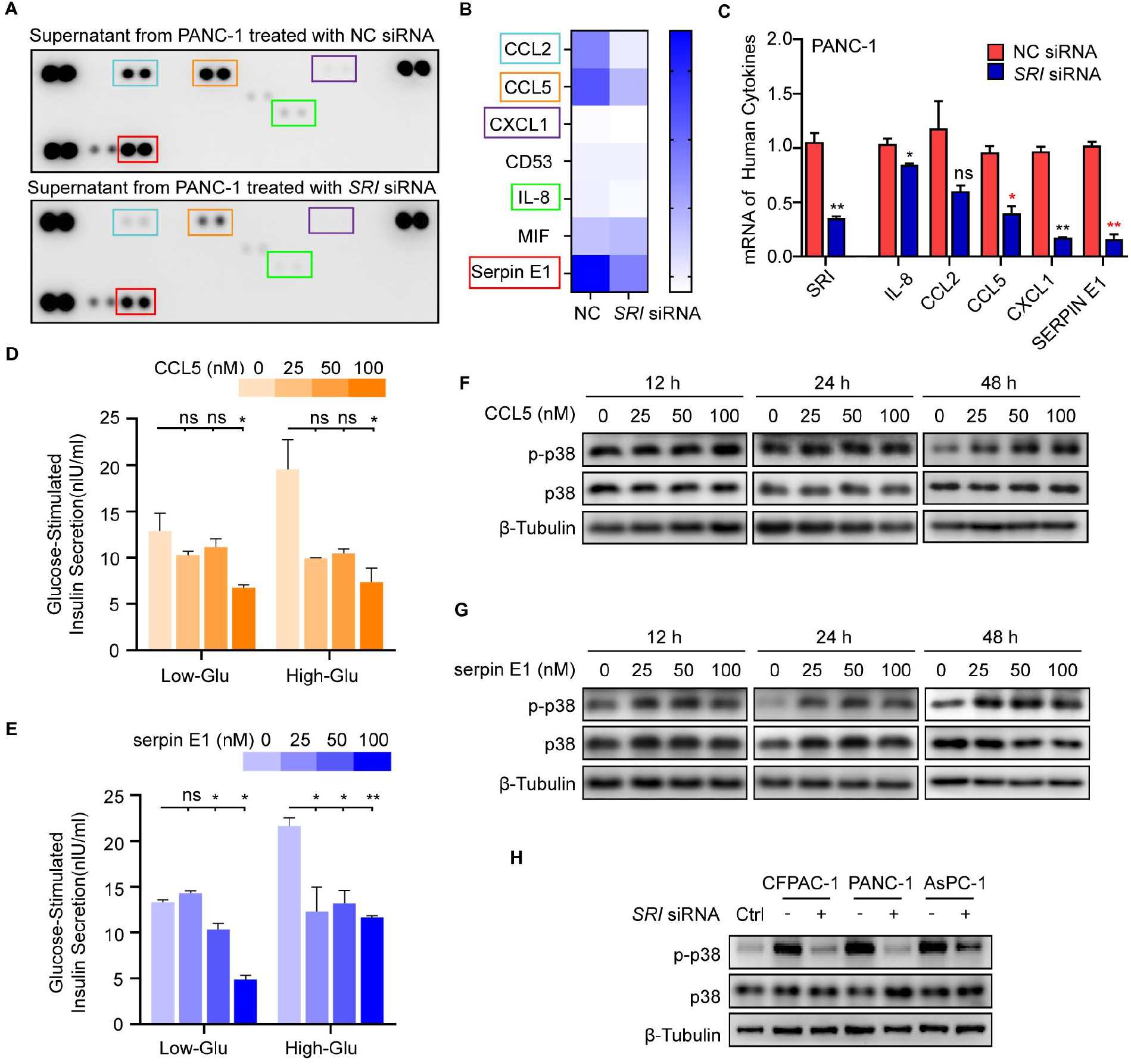
Sorcin-overexpressing PC cells release CCL5 and serpin E1 to inhibit insulin secretion in MIN6 cells. **(A)** Human cytokine array of supernatants from PANC-1 treated with CM-NC-siRNA and CM-*SRI*-siRNA. **(B)** Quantitative analysis of down-regulated components in PANC-1 treated with *SRI* siRNA in **A**. **(C)** Detection of *SRI* and 5 down-regulated cytokines’ mRNA by RT-PCR in PANC-1. The fresh culture media with inflammatory factors was changed every 12 hours until 72 hours after MIN6 cells were seeded and Insulin content in supernatant after GSIS in MIN6 cells treated with gradient concentrations of **(D)** CCL5 and **(E)** serpin E1. The phosphorylation levels of p38 in MIN6 cells incubated with different concentrations of **(F)** CCL5 and **(G)** serpin E1 at the concentration gradients of 0, 25, 50 and 100nM. **(H)** Detection of phosphorylation level of p38 pathway in MIN6 cells treated with different conditioned medium from PANC-1, CFPAC-1 and AsPC-1 cells for 72 hours. Ns, no significance; *P<0.05; **P<0.01, means ± SD was shown. Statistical analysis was performed by Student’s t-test analysis for two groups.

### Sorcin up-regulates CCL5 and serpin E1 expression by forming a positive feedback loop with STAT3

So far, we have shown that overexpression of sorcin in PC cells leads to enhanced secretion of CCL5 and serpin E1, which act on nearby β-cells. Since sorcin itself was not known to be a transcription factor, we speculated that it may interact with one or more transcription factors to up-regulate CCL5 and serpin E1 expression in PC cells. Indeed, sorcin has been reported to interact with STAT3 (Signal Transducer and Activator of Transcription 3) in mouse hepatocytes[43]. In PC cells, sorcin and STAT3 co-localize (Fig.5A) and can be co-immunoprecipitated as a protein complex (Fig. 5B-C). The phosphorylation level of STAT3 appeared to be dictated by the expression level of sorcin. In PANC-1 and AsPC-1 cells, phospho-STAT3 (p-STAT3) level increased with pcDNA-*SRI*-FLAG transfection in a concentration-dependent manner (Fig. 5D), and decreased with sorcin knockdown by siRNAs (Fig. 5E). Similarly, PC cell lines with elevated sorcin expression (Fig. 3B) had higher levels of p-STAT3 than the normal pancreatic duct epithelial cell line HPDE6 (Fig. 5F). In PDAC tumor tissues from human patients, sorcin was highly expressed in the cytoplasm of PC cells, while p-STAT3 as an activated transcription factor was enriched in the nucleus (Fig. S4A). Interestingly, when STAT3 expression was knocked down by siRNAs in PANC-1 cells, sorcin expression was also largely diminished (Fig. 5G), suggesting that STAT3 in turn can enhance the expression level of sorcin. Thus, the synergistic interactions between sorcin and STAT3 form a positive feedback loop (Fig. 5H), resulting in sustained overexpression of sorcin and activation of STAT3 in PC cells.

**Figure 5.**
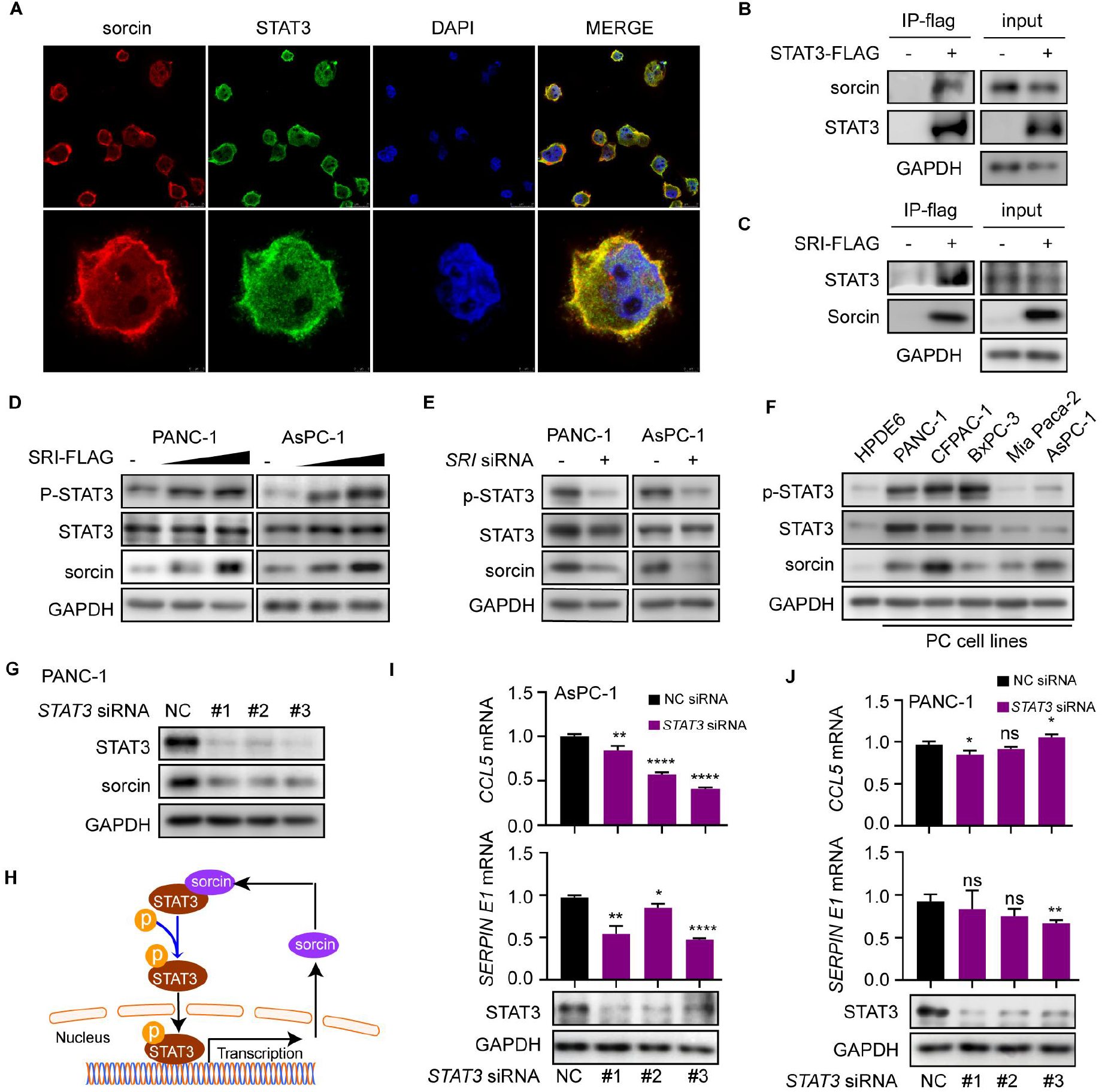
Sorcin up-regulates CCL5 and serpin E1 expression by forming a positive feedback loop with STAT3. **(A)** Confocal immunofluorescence microscopic images of sorcin (red) and STAT3 (green) in PANC-1 cells. **(B)** Immunoprecipitation of sorcin-STAT3 complex by anti-flag antibodies after pcDNA-*STAT3*-FLAG plasmid transfection in PANC-1 cells. **(C)** Immunoprecipitation of sorcin-STAT3 complex by anti-flag antibodies after pcDNA-*SRI*-FLAG transfection in PANC-1 cells. **(D)** The phosphorylation levels of STAT3 in PANC-1 and AsPC-1 transfected with gradient increasing pcDNA-*SRI*-FLAG plasmids at different concentration from 1 to 2 μg/well in 12-well plates, taking empty pcDNA vector as control. **(E)** The phosphorylation levels of STAT3 in PANC-1 and AsPC-1 transfected with *SRI* siRNA. **(F)** The levels of sorcin and STAT3, and the phosphorylation levels of STAT3 in HPDE6 cell and 5 kinds of PC cells. **(G)** The level of sorcin in PANC-1 transfected with *STAT3* siRNA. **(H)** A Scientific hypothesis diagram of sorcin-STAT3 positive feedback loop. Detection of *CCL5* and *SERPIN E1* mRNA, and the expression levels of STAT3 after treating with *STAT3* siRNA in **(I)** AsPC-1 and **(J)** PANC-1. Ns, no significance; *P<0.05; **P<0.01; ***P<0.001; ****P<0.0001, means ± SD was shown. Statistical analysis was performed by Student’s t-test analysis for two groups.

To further confirm that the sorcin-STAT3 loop is responsible for up-regulating the transcription of CCL5 and serpin E1 in PC cells, we examined the effect of STAT3 knockdown on *CCL5* and *SERPIN E1* transcript levels. In AsPC-1 and CFPAC-1 cells, STAT3 knockdown via three different siRNAs all resulted in down-regulation of *CCL5* and *SERPIN E1* transcripts (Fig. 5J and Supplementary Fig. S4B), similar to what we observed with sorcin knockdown (Fig. 4C and Fig. S4A-D). In PANC-1 cells, on the other hand, the impact of STAT3 knockdown on *CCL5* mRNA level varied with different siRNAs, while all three STAT3-siRNAs led to a slight but significant decrease in *SERPIN E1* mRNA. The discrepancy is perhaps not surprising, considering that STAT3 knockdown not only affects gene targets directly downstream of the sorcin-STAT3 loop (Fig. 5J), but also disrupts the interactions between STAT3 and other proteins[44].

### *SRI* can be used to differentiate PCAND from T2DM and downstream serpin E1 may be the potential biomarker

For validation with large-scale cohorts, a total of 13136 T2DM and 102 PCAND were identified in UK Biobank (Fig. 6A). With a unified clinical model (Materials and Methods), 4 independent models (#1 Clinical risk model, #2 “clinical+*SRI*”, #3 “clinical+*CDKN2A*”, #4 “clinical+*KRAS*”) and a combined model (#5 “clinical+*SRI*+*CDKN2A*+*KRAS*”) were developed and then validated in the testing set. Among them, the *SRI*-based model (#2 “clinical+*SRI*”) outperformed other driver genes with higher AUROC of 0.768 (95% CI 0.553-0.896) (Fig. 6B and Supplementary Table 3), which further support the notion that early screening strategies based on *SRI* may be better than other oncogenes. Moreover, when *SRI*, *CDKN2A* and *KRAS* were combined, the model (#5 “clinical+*SRI*+*CDKN2A*+*KRAS*”) achieved the highest AUROC of 0.772 (95% CI 0.578-0.912) (Fig. 6B and Supplementary Table 3), which suggested that future studies should focus on the synergistic effect of *SRI* and other oncogenes.

**Figure 6.**
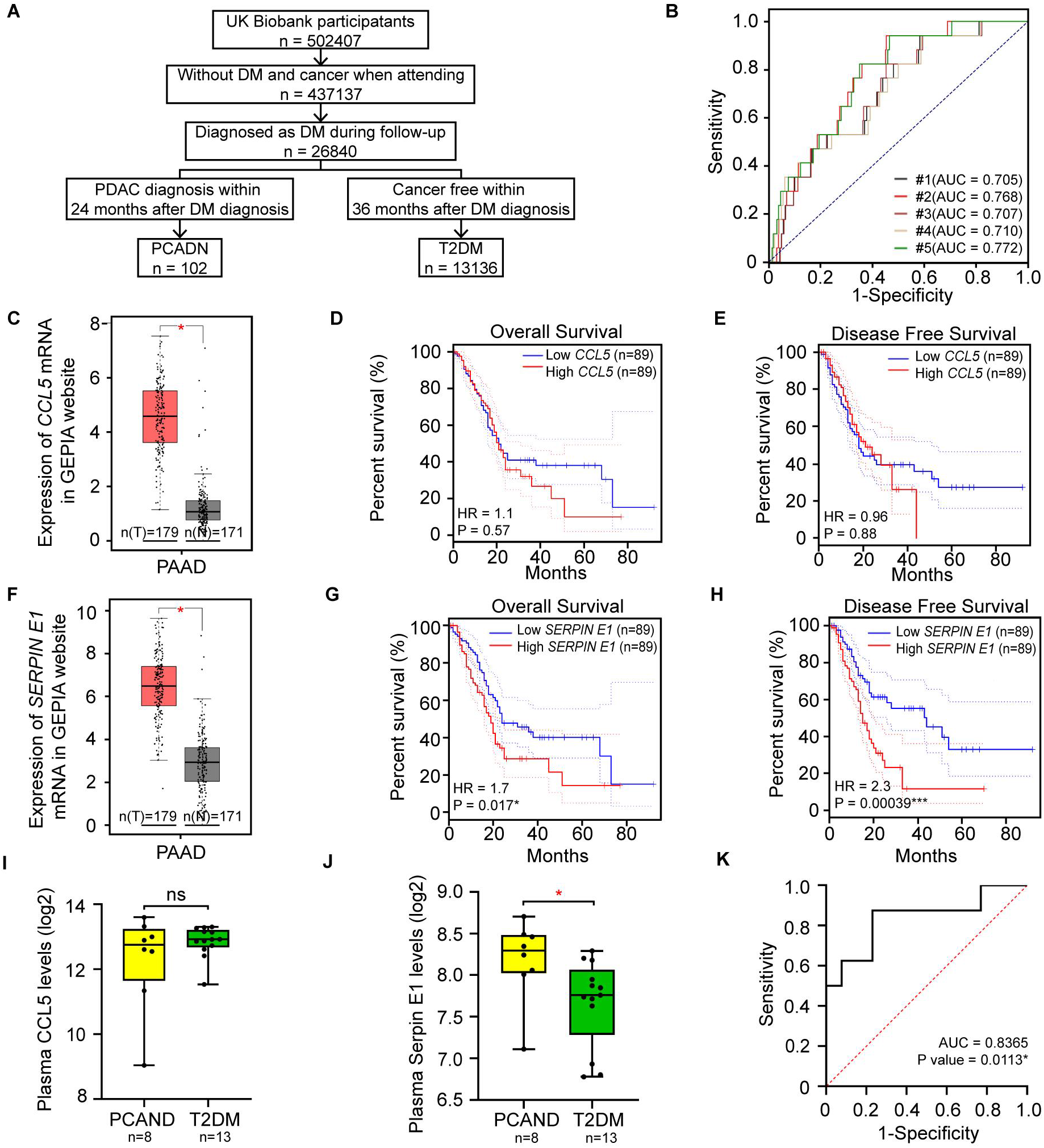
*SRI* can be used to differentiate PCAND from T2DM and downstream Serpin E1 may be the potential biomarker. **(A)** The procedure of population screening and partitioning into PCAND and pure T2DM cohorts in the UK Biobank. **(B)** Evaluation performance of five models in the testing set. #1 Clinical risk model, #2 clinical+*SRI*, #3 clinical+*CDKN2A*, #4 “clinical+*KRAS*, #5 clinical+*SRI*+*CDKN2A*+*KRAS*. **(C)** The expression pattern of *CCL5* mRNA in PC tumor (T) compare with normal pancreas tissue (N). Comparison of **(D)** overall survival rates and **(E)** disease free survival rates between high *CCL5* group and low *CCL5* group, taking the median of *CCL5* expression as the cut-off. **(F)** The expression pattern of *SERPIN E1* mRNA in PC tumor (T) compare with normal pancreas tissue (N). Comparison of **(G)** overall survival rates and **(H)** disease free survival rates between high *SERPIN E1* group and low *SERPIN E1* group, taking the median of *SERPIN E1* expression as the cut-off. **C-H** are based on The Cancer Genome Atlas (TCGA) database and Genotype-Tissue Expression (GTEx) from GEPIA website. The plasma **(I)** CCL5 and **(J)** serpin E1 levels in pancreatic cancer associated new-onset diabetes (PCAND) and type 2 diabetes (T2DM). **(K)** The sensitivity and specificity in discriminating PCAND patients from DM population for serpin E1 are shown. The ROC curves and AUC were calculated in 8 PCADN patients and 13 pure T2DM patients. Ns, no significance; *P<0.05; ***P<0.001, means ± SD was shown. Statistical analysis was performed by Student’s t-test analysis for two groups. And Kaplan-Meier curves of survival were compared using the log-rank test.

According to TCGA and GTEx data, both *CCL5* and *SERPIN E1* were expressed at significantly higher levels in pancreatic cancer tissues (T, n = 179) than in nearby normal pancreatic tissues (N, n = 171) (Fig. 6C and 6F). Among PAAD patients, a high *SERPIN E1* expression was associated with worse prognosis of pancreatic cancer patients, while the differences between high *CCL5* and low *CCL5* groups did not reach statistical significance (Fig. 6D-E and 6G-H).

To assess the performance of CCL5 and serpin E1 as potential biomarkers for PCAND, we measured their concentrations in peripheral blood from patients diagnosed with either PCAND (n = 8) or T2DM (n = 13). There was no difference in CCL5 between the two groups (Fig. 6I), but the level of serpin E1 was significantly higher in PCAND than in pure T2DM (Fig. 6J). In this small cohort, serpin E1 achieved an AUROC of 0.8364 in differentiating between PCAND and pure T2DM (*P* = 0.0113, 95% CI 0.6415-1.000) (Fig. 6K), demonstrating its potential utility as a biomarker for PCAND.

## DISCUSSION

Pancreatic cancer (PC), a devastating disease characterized by late diagnosis, limited treatment success and dismal prognosis, remains a major medical challenge. A rise in blood glucose is one of the early warning signs of underlying PC, which could be the externalization of genetic events in PC progression. Considering the convenience and popularity of blood glucose monitoring, one of the key to early diagnosis of PC is to identify the only 1% PCAND in new-onset diabetes population as early as possible[45-47]. To make progress on PCAND detection requires an improved understanding of the molecular mechanisms and signaling pathways underlying its specific pathogenesis.

In this study, we have mapped out a novel sorcin-STAT3-Serpin E1/CCL5 signaling axis in PC cells, which explains how early pre-symptomatic PC may coincide with new-onset diabetes in some patients[48]. In PC cells cultured *in vitro*, sorcin and STAT3 form a positive feedback loop to enhance the transcription of serpin E1 and CCL5. These inflammatory cytokines released by PC cells, can impair nearby islet β-cells, likely by activating the p38 signaling pathway. On top of that, in biopsies obtained from 88 PDAC patients, we found an elevated expression of sorcin in pancreatic cancer tissues, especially so in PCAND. These results suggest that sorcin could be the key driver in PCAND and aberrant activation of the sorcin-STAT3-Serpin E1/CCL5 signaling axis likely underlie PCAND pathogenesis.

During exploring the driver mechanism of *SRI* in PCADN, we have figured out a potential bidirectional crosstalk between PCAND pathogenesis and inflammation, likely regulating by sorcin-STAT3-Serpin E1/CCL5 signaling axis. Interestingly, the signaling axis we describe here shares a common critical node, STAT3, with the inflammatory pathway downstream of KRAS, whose mutations are the most common genetic abnormality in PC[6]. Given the long time (over 10 years for PC development)[49] and inadequacy of *KRAS* mutations to PC progression[8], it remains far from reaching clinics for screening or diagnostic use. In this study, we found fasting blood glucose level in pure PC and PCAND patients before pancreatectomy was positively correlated with sorcin expression level. Therefore, the rise in blood glucose driven by *SRI* gene could be the externalization of PCAND, typically manifesting 2-3 years prior to the diagnosis of PC[25], most likely during the progression from PanIN-3 to PC (figure 7). These results further support the notion that early screening strategies based on *SRI* gene may be better than *KRAS* and other oncogenes that are mutated in early PanINs stage.

**Figure 7.**
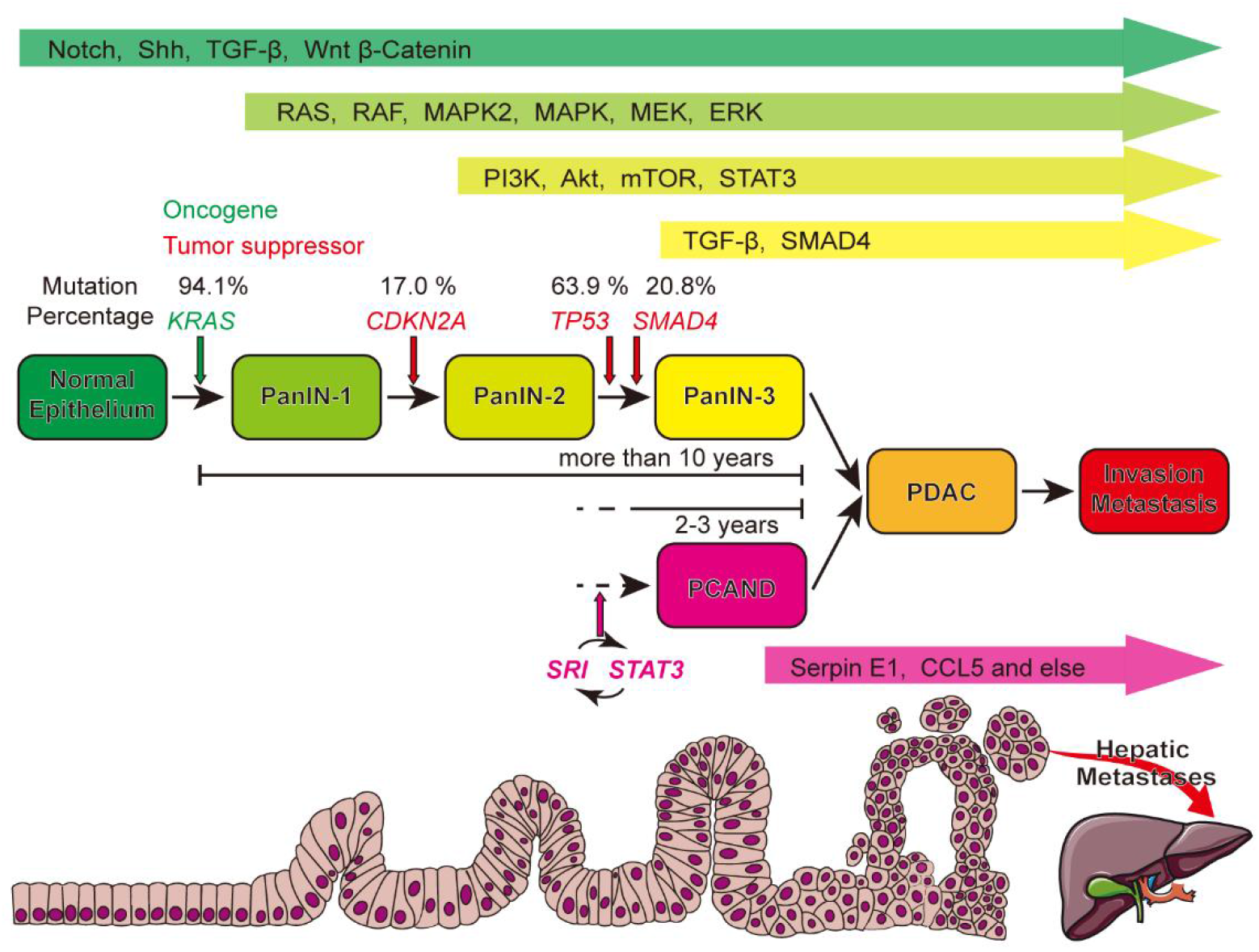
*SRI* can be the driver gene in the pathogenesis of PCAND. During PC genesis and development, a variety of driver gene mutation (*KRAS*, *CDKN2A*, *TP53* and *SMAD4*) and signaling pathways deregulation participate in the progression from pancreatic intraepithelial neoplastic lesions (PanINs 1-3) to PDAC. *KRAS* mutation is the initiating genetic event for PC and the progression of normal pancreatic tissue to PDAC involves a stepwise genetic transition projected to span more than 10 years. And PCAND was observed new-onset diabetes within 2–3 years prior of PDAC, so abnormal activation of the sorcin-STAT3 positive feedback loop may occur during the progression from PanIN 1-3 to PDAC, later than oncogene mutation. And The downstream inflammatory cytokines/chemokines (like CCL5 and serpin E1) may be potential biomarkers. (Compiled from data in Jones et al. 2008, Biankin et al. 2012, Sonja Maria Wörmann et al. 2013, Sausen et al. 2015, Waddell et al. 2015, Witkiewicz et al. 2015, Mohammad Aslam Aslam Khan et al. 2017 and Andrew M Waters et al. 2018)

Previous research suggested that sorcin acted as a protective factor to β-cells in T2DM[27]. However, in our *in vitro* research, PC-derived sorcin play a negative role in β-cells function and can induce an inflammatory damage. Unlike T2DM with adipocyte-derived inflammatory cytokines [50], PC has specific inflammatory tumor microenvironment(TME)[51]. In this study, the increased secretion of serpin E1 and CCL5 induced by sorcin-STAT3 interaction may in turn contribute to the formation of an inflammatory TME[52], alongside *KRAS* associated inflammatory signaling[53]. Further, based on large-scale cohorts from UK Biobank, we confirmed that *SRI*-based model is superior to those models based on other driver genes like *KRAS* in differentiating PCAND from T2DM, and the combination of *SRI*, *KRAS*, *CDKN2A* with clinical model can further increase the efficiency. On the other hand, based on a small cohort of PCAND and T2DM patients, the concentration of serpin E1 in peripheral blood samples showed decent diagnostic performance. We are aware that it is a preliminary study, and still has several limitations, such as the sample size of the clinical cohorts. As a next step, we believe that a larger-scale validation study with a longitudinal sampling scheme should be carried out in the future.

In summary, our study identified a novel sorcin-STAT3-Serpin E1/CCL5 signaling axis as a key driver of PCAND pathogenesis. The convergence of sorcin and KRAS signaling on STAT3 suggests a potential bidirectional crosstalk, which should be factored in when considering targeted therapies for PC involving these pathways. Our results also suggest that clinical-*SRI* gene combination may be the new strategy to identify PCAND and serpin E1 may be a potential biomarker for PCAND. Further studies on the molecules downstream of the sorcin pathway may yield valuable clues for the early diagnosis of PC.

## Supporting information

All Supplemental Figures and Tables

## References

1. Mizrahi, J.D., et al., Pancreatic cancer. Lancet (London, England), 2020. 395(10242): p. 2008–2020.

2. Gressens, S.B., et al., Controversial Roles of the Renin Angiotensin System and Its Modulators During the COVID-19 Pandemic. Frontiers in physiology vol., (1664-042X (Print)).

3. Poruk, K.E., et al., Screening for pancreatic cancer: why, how, and who? Annals of Surgery, 2013. 257(1): p. 17–26.

4. Kanda, M., et al., Presence of somatic mutations in most early-stage pancreatic intraepithelial neoplasia. Gastroenterology, 2012. 142(4).

5. Waddell, N., et al., Whole genomes redefine the mutational landscape of pancreatic cancer. Nature, 2015. 518(7540): p. 495–501.

6. Makohon-Moore, A. and C.A. Iacobuzio-Donahue, Pancreatic cancer biology and genetics from an evolutionary perspective. Nature Reviews. Cancer, 2016. 16(9): p. 553–565.

7. Waters, A.M. and C.J. Der, KRAS: The Critical Driver and Therapeutic Target for Pancreatic Cancer. Cold Spring Harbor Perspectives In Medicine, 2018. 8(9).

8. Siveke, J.T., et al., Concomitant pancreatic activation of Kras(G12D) and Tgfa results in cystic papillary neoplasms reminiscent of human IPMN. Cancer Cell, 2007. 12(3): p. 266–279.

9. Sharma, A., et al., Fasting Blood Glucose Levels Provide Estimate of Duration and Progression of Pancreatic Cancer Before Diagnosis. Gastroenterology, 2018. 155(2): p. 490–500.e2.

10. Bao, J., et al., Pancreatic cancer-associated diabetes mellitus is characterized by reduced β-cell secretory capacity, rather than insulin resistance. Diabetes Research and Clinical Practice, 2022. 185: p. 109223.

11. Permert, J., et al., Improved glucose metabolism after subtotal pancreatectomy for pancreatic cancer. British Journal of Surgery, 1993. 80(8): p. 1047–1050.

12. Huang, H., et al., Novel Blood Biomarkers of Pancreatic Cancer–Associated Diabetes Mellitus Identified by Peripheral Blood–Based Gene Expression Profiles. Official journal of the American College of Gastroenterology | ACG, 2010. 105(7).

13. Kang, M., et al., VNN1, a potential biomarker for pancreatic cancer-associated new-onset diabetes, aggravates paraneoplastic islet dysfunction by increasing oxidative stress. Cancer letters, 2016. 373(2): p. 241–250.

14. Pfeffer, F., et al., Expression of Connexin26 in Islets of Langerhans Is Associated With Impaired Glucose Tolerance in Patients With Pancreatic Adenocarcinoma. Pancreas, 2004. 29(4).

15. Liao, W.-C., et al., Galectin-3 and S100A9: Novel Diabetogenic Factors Mediating Pancreatic Cancer– Associated Diabetes. Diabetes Care, 2019. 42(9): p. 1752.

16. Basso, D., et al., Pancreatic cancer-derived S-100A8 N-terminal peptide: A diabetes cause? Clinica Chimica Acta, 2006. 372(1): p. 120–128.

17. Ding, X., et al., Pancreatic cancer cells selectively stimulate islet beta cells to secrete amylin. Gastroenterology, 1998.

18. Kolb, A., et al., Glucagon/insulin ratio as a potential biomarker for pancreatic cancer in patients with new-onset diabetes mellitus. Cancer Biology & Therapy, 2009. 8(16): p. 1527–1533.

19. Krechler, T., et al., Polymorphism -23HPhI in the promoter of insulin gene and pancreatic cancer: a pilot study. Neoplasma, 2009.

20. Javeed, N., et al., Pancreatic Cancer-Derived Exosomes Cause Paraneoplastic β-cell Dysfunction. Clinical cancer research : an official journal of the American Association for Cancer Research, 2015. 21(7): p. 1722–1733.

21. Permert, J., et al., Islet Amyloid Polypeptide in Patients with Pancreatic Cancer and Diabetes. New England Journal of Medicine, 1994. 330(5): p. 313–318.

22. Sharaf, R.N., et al., Computational prediction and experimental validation associating FABP-1 and pancreatic adenocarcinoma with diabetes. BMC gastroenterology, 2011. 11: p. 5–5.

23. Basso, D., et al., Insulin-like growth factor-I, interleukin-1 alpha and beta in pancreatic cancer: role in tumor invasiveness and associated diabetes. Int J Clin Lab Res, 1995.

24. Abbruzzese, J.L., et al., The Interface of Pancreatic Cancer With Diabetes, Obesity, and Inflammation: Research Gaps and Opportunities: Summary of a National Institute of Diabetes and Digestive and Kidney Diseases Workshop. Pancreas, 2018. 47(5): p. 516–525.

25. Chari, S.T., et al., Pancreatic cancer-associated diabetes mellitus: prevalence and temporal association with diagnosis of cancer. Gastroenterology, 2008. 134(1).

26. Sharma, A., et al., Model to Determine Risk of Pancreatic Cancer in Patients With New-Onset Diabetes. Gastroenterology, 2018. 155(3).

27. Marmugi, A., et al., Sorcin Links Pancreatic β-Cell Lipotoxicity to ER Ca2+ Stores. Diabetes, 2016. 65(4): p. 1009–1021.

28. Sharma, S., et al., Predicting Pancreatic Cancer in the UK Biobank Cohort Using Polygenic Risk Scores and Diabetes Mellitus. Gastroenterology, 2022. 162(6).

29. Diagnosis and classification of diabetes mellitus. Diabetes Care, 2013. 36 Suppl 1: p. S67–S74.

30. UK Biobank data on 500,000 people paves way to precision medicine. Nature, 2018. 562(7726): p. 163–164.

31. Sah, R.P., et al., New insights into pancreatic cancer-induced paraneoplastic diabetes. Nature Reviews. Gastroenterology & Hepatology, 2013. 10(7): p. 423–433.

32. Perera, C.J., et al., Role of Pancreatic Stellate Cell-Derived Exosomes in Pancreatic Cancer-Related Diabetes: A Novel Hypothesis. Cancers, 2021. 13(20).

33. Pang, W., et al., Pancreatic cancer-derived exosomal microRNA-19a induces β-cell dysfunction by targeting ADCY1 and EPAC2. International journal of biological sciences, 2021. 17(13): p. 3622–3633.

34. Talchai, C., et al., Pancreatic β cell dedifferentiation as a mechanism of diabetic β cell failure. Cell, 2012. 150(6): p. 1223–1234.

35. Zhang, C., et al., MafA is a key regulator of glucose-stimulated insulin secretion. Molecular and cellular biology, 2005. 25(12): p. 4969–4976.

36. Zhao, L., et al., The islet beta cell-enriched MafA activator is a key regulator of insulin gene transcription. J Biol Chem, 2005.

37. Fujimoto, K. and K.S. Polonsky, Pdx1 and other factors that regulate pancreatic beta-cell survival. Diabetes, obesity & metabolism, 2009. 11 Suppl 4(Suppl 4): p. 30–37.

38. Chandra, V., et al., RFX6 regulates insulin secretion by modulating Ca2+ homeostasis in human β cells. Cell Rep, 2014.

39. Greer, J.B. and D.C. Whitcomb, Inflammation and pancreatic cancer: an evidence-based review. Current Opinion In Pharmacology, 2009. 9(4): p. 411–418.

40. Padoan, A., M. Plebani, and D. Basso, Inflammation and Pancreatic Cancer: Focus on Metabolism, Cytokines, and Immunity. International Journal of Molecular Sciences, 2019. 20(3).

41. Donath, M.Y., et al., Mechanisms of β-Cell Death in Type 2 Diabetes. Diabetes, 2005. 54(suppl 2): p. S108.

42. Wei, X., et al., Inhibition of p38 mitogen-activated protein kinase exerts a hypoglycemic effect by improving β cell function via inhibition of β cell apoptosis in db/db mice. Journal of enzyme inhibition and medicinal chemistry, 2018. 33(1): p. 1494–1500.

43. Li, X., et al., Negative Regulation of Hepatic Inflammation by the Soluble Resistance-Related Calcium-Binding Protein Signal Transducer and Activator of Transcription 3. Frontiers In Immunology, 2017. 8: p. 709.

44. Zou, S., et al., Targeting STAT3 in Cancer Immunotherapy. Molecular Cancer, 2020. 19(1): p. 145.

45. Singhi, A.D., et al., Early Detection of Pancreatic Cancer: Opportunities and Challenges. Gastroenterology, 2019. 156(7): p. 2024–2040.

46. Chari, S.T., et al., Probability of pancreatic cancer following diabetes: a population-based study. Gastroenterology, 2005. 129(2): p. 504–511.

47. Owens, D.K., et al., Screening for Pancreatic Cancer: US Preventive Services Task Force Reaffirmation Recommendation Statement. JAMA, 2019. 322(5): p. 438–444.

48. Chung, H.H., K.S. Lim, and J.K. Park, Clinical Clues of Pre-Symptomatic Pancreatic Ductal Adenocarcinoma Prior to Its Diagnosis: A Retrospective Review of CT Scans and Laboratory Tests. Clinics and Practice, 2022. 12(1): p. 70–77.

49. Yachida, S., et al., Distant metastasis occurs late during the genetic evolution of pancreatic cancer. Nature, 2010. 467(7319): p. 1114–1117.

50. Bruun, J.M., et al., Regulation of adiponectin by adipose tissue-derived cytokines: in vivo and in vitro investigations in humans. American Journal of Physiology. Endocrinology and Metabolism, 2003. 285(3): p. E527–E533.

51. Ho, W.J., E.M. Jaffee, and L. Zheng, The tumour microenvironment in pancreatic cancer - clinical challenges and opportunities. Nature Reviews. Clinical Oncology, 2020. 17(9): p. 527–540.

52. Aldinucci, D., C. Borghese, and N. Casagrande, The CCL5/CCR5 Axis in Cancer Progression. Cancers, 2020. 12(7).

53. Kitajima, S., R. Thummalapalli, and D.A. Barbie, Inflammation as a driver and vulnerability of KRAS mediated oncogenesis. Seminars In Cell & Developmental Biology, 2016. 58: p. 127–135.

