## Supplementary material for "Sorcin-STAT3-Serpin E1/CCL5 axis can be the trigger of pancreatic cancer-associated new-onset diabetes": All Supplemental Figures and Tables

**SUPPLEMENTARY FILES**

**Supplementary Table 1. Antibodies used**

| Name | Company | Dilution | |
|---|---|---|---|
| Sorcin | Proteintech | WB 1:1000 | IF 1:100 |
| Sorcin | Abcam | IHC 1:500 | |
| STAT3 | Proteintech | WB 1:1000 | IF 1:100 |
| p-STAT3 | Proteintech | WB 1:1000 | |
| p-STAT3 | CST | IHC 1:500 | |
| Insulin | Proteintech | IF 1:100 | |
| Insulin | Abcam | WB 1:1000 | IHC 1:68000 |
| GAPDH | Abcam | WB 1:10000 | |
| p38 | HUABIO | WB 1:1000 | |
| p-p38 | HUABIO | WB 1:1000 | |
| Actin | Proteintech | WB 1:10000 | |
| Tubulin | Proteintech | WB 1:10000 | |

**Supplementary Table 2. Primer sequences used**

| Name | Primer Sequence (Forward 5' - 3') | Primer Sequence (Reverse 5' - 3') |
|---|---|---|
| *GAPDH (human)* | GCACCGTCAAGGCTGAGAAC | TGGTGAAGACGCCAGTGGA |
| *STAT3 (human)* | AACTCTCACGGACGAGGAGCT | AGTAGTGAACTGGACGCCGG |
| *SRI (human)* | TAAAGAACTCTGGGCTGTACTG | GTGCTGTATCGTTTTGCAATTG |
| *IL8(human)* | AACTGAGAGTGATTGAGAGTGG | ATGAATTCTCAGCCCTCTTCAA |
| *CCL2(human)* | ATGACTTCCAAGCTGGCCGTGGCT | ATGACTTCCAAGCTGGCCGTGGCT |
| *CCL5(human)* | GGCAGCCCTCGCTGTCATCCTCA | CTTGATGTGGGCACCGGGGCAGTG |
| *SERPIN E1(human)* | AACGTGGTTTTCTCACCCTAT | CAATCTTGAATCCCATAGCTGC |
| *CXCL1(human)* | AAGAACATCCAAAGTGTGAACG | CACTGTTCAGCATCTTTTCGAT |
| *Gadph (mouse)* | GGTGAAGGTCGGAGTCAACG | CAAAGTTGTCATGGATGHACC |
| *Ins (mouse)* | TTGTCAAACAGCATCTTTGTGG | GGACTTGGGTGTGTAGAAGAAG |
| *Pdx1 (mouse)* | GACATCTCCCCATACGAAGTG | GTGAGCTTTGGTGGATTTCATC |
| *Mafa (mouse)* | GCTTCAGCAAGGAGGAGGTCATC | GGCACTTCTCGCTCTCCAGAATG |
| *Foxo1 (mouse)* | AATTCACCCAGTCCAAACTACT | TACTGGTTCAATCCTCCGTAAC |
| *Rfx6 (mouse)* | ACATCTGCCATGAACCGCTTGAC | CACCCATCCTACCATAGCCATTGC |

**Supplementary Table 3. Evaluation performance of five models in the testing set**

| No | Model | AUC (95% CI) | Sensitivity | Specificity | Accuracy |
|---|---|---|---|---|---|
| 1 | Clinical | 0.705(0.523-0.883) | 0.941 | 0.417 | 0.421 |
| 2 | Clinical + *SRI* | 0.768(0.553-0.896) | 0.941 | 0.546 | 0.550 |
| 3 | Clinical + *CDKN2A* | 0.707(0.474-0.905) | 0.824 | 0.547 | 0.550 |
| 4 | Clinical + *KRAS* | 0.710(0.451-0.886) | 0.941 | 0.412 | 0.416 |
| 5 | Clinical+*SRI*+ *CDKN2A*+*KRAS* | 0.772(0.578-0.912) | 0.941 | 0.535 | 0.538 |

**Supplementary Figure 1**

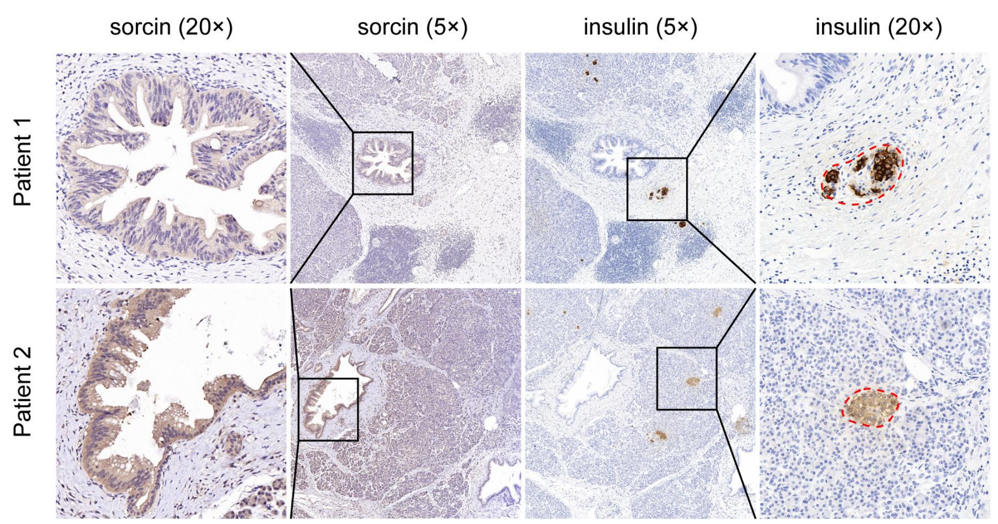


**Figure S1. SRI is highly expressed in pancreatic cancer-associated new-onset diabetes**

Negative correlation between the IHC intensity of sorcin in pancreatic cancer cells and IHC intensity of insulin in adjacent pancreatic islets.

**Supplementary Figure 2**

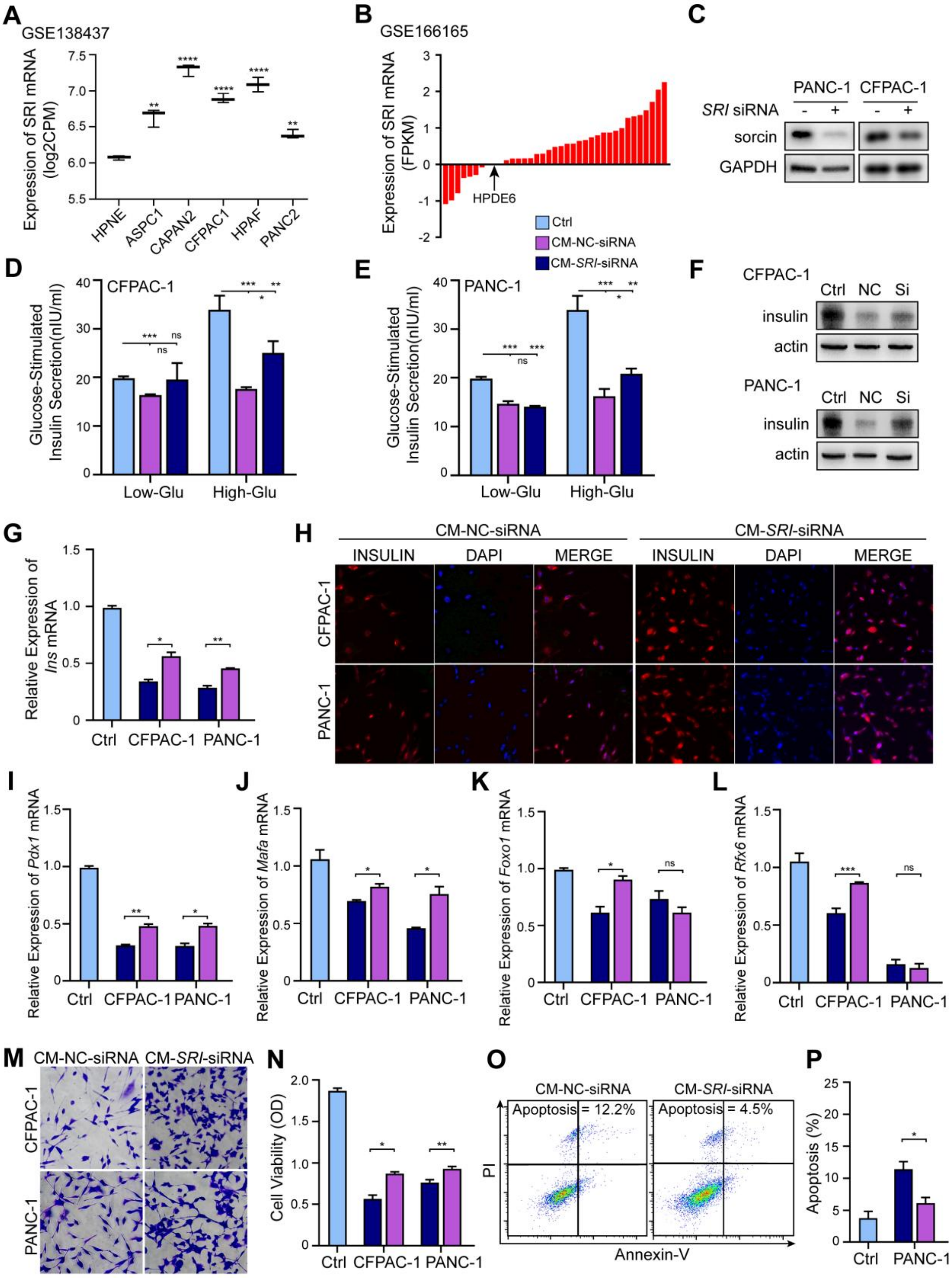

**Figure S2. PC cells inhibit insulin secretion in MIN6 cells in a sorcin-dependent manner**

*SRI* expression in PC cell lines and normal pancreatic duct cell line (HPNE/HPDE6) based on **(A)** GSE138437 dataset and **(B)** the GSE166165 dataset. **(C)** Detection of knockdown effect in PANC-1 and CFPAC-1 cells by immunoblotting. Insulin content in supernatant after GSIS in MIN6 cells incubated with conditioned medium form **(D)** PANC-1 and **(E)** CFPAC-1 cells pretreated with NC siRNA (CM-NC-siRNA) and SRI siRNA (CM-*SRI*-siRNA). **(F)** Detection of insulin content in MIN6 cells incubated with CM-NC-siRNA and CM-*SRI*-siRNA from PANC-1 and CFPAC-1 cells by immunoblotting. **(G)** The expression of *Ins* mRNA in MIN6 cells incubated with different conditioned medium from PANC-1 and CFPAC-1 cells. **(H)** Immunofluorescence shows the content of insulin in MIN6 cells treated by different conditioned medium from PANC-1 and CFPAC-1 cells. The expression of **(I)** *Pdx1*, **(J)** *Mafa*, **(K)** *Foxo1* and **(L)** *Rfx6* mRNA in MIN6 cells incubated with different conditioned medium from PANC-1 and CFPAC-1 cells. **(M)** Morphology of MIN6 cells treated with different conditioned medium from PANC-1 and CFPAC-1 cells. **(N)** Detection of cell viability of MIN6 cells treated with different conditioned medium from PANC-1 and CFPAC-1 cells by MTT assays. **(O)** Detection of apoptosis by flow cytometry in MIN6 treated with different conditioned medium from PANC-1. **(P)** Quantification results of apoptosis rate. Ns, no significance; *P<0.05; **P<0.01; ***P<0.001; ****P<0.0001, means ± SD was shown. Statistical analysis was performed by Student's t-test analysis for two groups.

**Supplementary Figure 3**

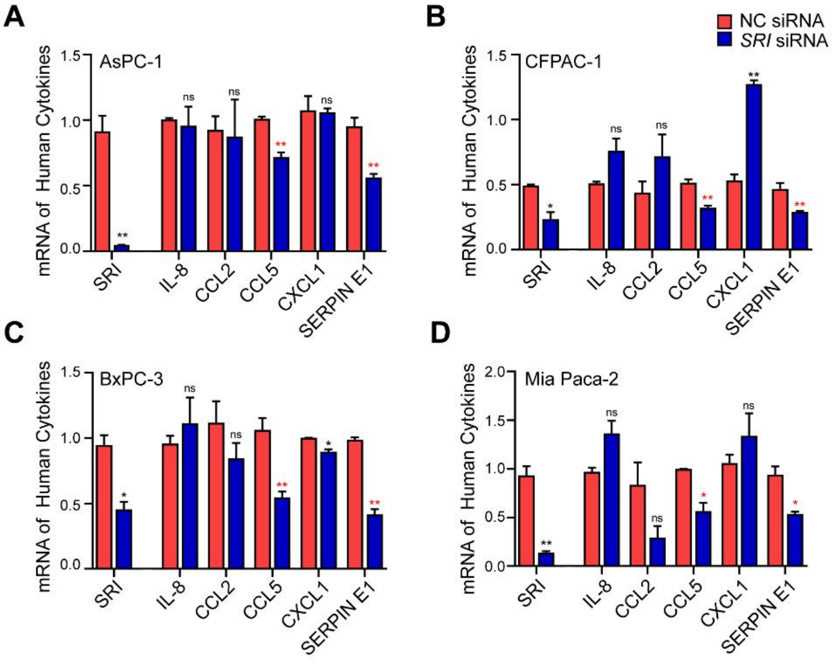


**Figure S3. Sorcin-overexpressing PC cells release CCL5 and serpin E1 to inhibit insulin secretion in MIN6 cells**

Detection of *SRI* and 5 down-regulated cytokines' mRNA by RT-PCR in **(A)** AsPC-1, **(B)** CFPAC-1, **(C)** BxPC-3 and **(D)** Mia Paca-2. Ns, no significance; *P<0.05; **P<0.01, means ± SD was shown. Statistical analysis was performed by Student's t-test analysis for two groups.

**Supplementary Figure 4**

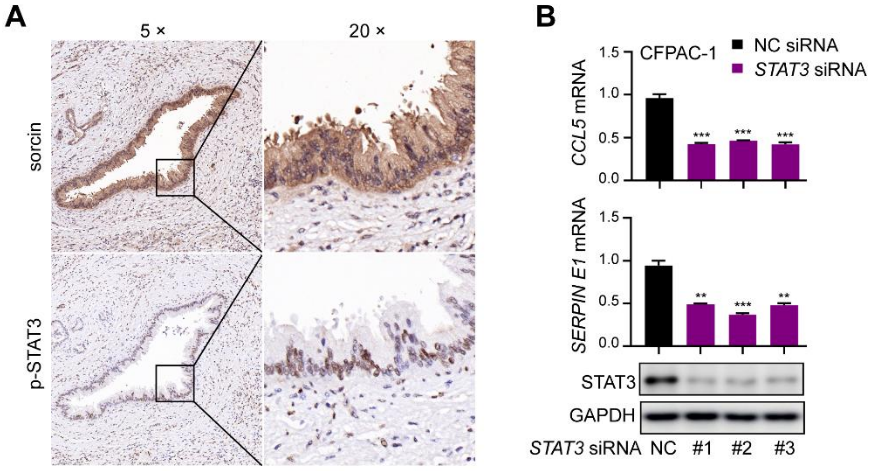


**Figure S4. Sorcin up-regulates CCL5 and serpin E1 expression by forming a positive feedback loop with STAT3**

**(A)** Immunohistochemistry staining shown that sorcin was highly expressed in cytoplasm, and p-STAT3 was aggregated in the nucleus in patients with pancreatic cancer. **(B)** Detection of *CCL5* and *SERPIN E1* mRNA, and the expression levels of STAT3 after treating with *STAT3* siRNA in CFPAC-1. **P<0.01; ***P<0.001, means ± SD was shown. Statistical analysis was performed by Student's t-test analysis for two groups.
